# Selfing species exhibit diminished niche breadth over time

**DOI:** 10.1101/157974

**Authors:** Daniel S. Park, Aaron M. Ellison, Charles C. Davis

**Author notes:** Correspondence should be addressed to: Daniel S. Park Department of Organismic and Evolutionary Biology, 22 Divinity Ave. Harvard University Herbaria, Cambridge, MA 02138.

## Abstract

Self-pollinating plants (“selfers”) have larger geographic ranges and inhabit higher latitudes than their outcrossing relatives. This finding has led to the hypothesis that selfers also have broader climatic niches. It is possible that the increased likelihood of successful colonization into new areas and the initial purging of deleterious mutations may offset selfers’ inability to adapt to new environments due to low heterozygosity. Here, for the first time, we examine the climatic niches and mutation accumulation rates of hundreds of closely related selfing and outcrossing species. Contrary to expectations, selfers do not have wider climatic niche breadths than their outcrossing sister taxa despite selfers’ greatly expanded geographic ranges. Selfing sister pairs also exhibit greater niche overlap than outcrossing sisters, implying that climatic niche expansion becomes limited following the transition to selfing. Further, the niche breadth of selfers is predicted to decrease significantly faster than that of closely-related outcrossers. In support of these findings, selfers also display significantly higher mutation accumulation rates than their outcrossing sisters, implying decreased heterozygosity, effective population size, and adaptive potential. These results collectively suggest that while the release from mate limitation among selfing species may result in initial range expansion, range size and niche breadth are decoupled, and the limitations of an increasingly homogeneous genome will constrict selfers’ climatic niches and over time reduce their geographic ranges.

## Introduction

Species ranges are influenced by various life-history traits (Sexton *et al.* 2009), including the evolution of autonomous reproduction (Igic & Busch 2013). “Baker’s Law” posits that the ability of a species to self-fertilize increases colonization and establishment success by bypassing mate limitation and pollinator requirements (Baker 1955; Stebbins 1957; Pannell & Barrett 1998). Along these lines, self-pollinating plant species (“selfers”) consistently display larger geographic ranges and occupy higher maximum latitudes than closely related outcrossing species (Grossenbacher *et al.* 2015). Based on these results, it has been hypothesized that selfing species may have greater climatic tolerances and climatic niche breadths (hereafter “niche breadths”) than outcrossing sister taxa (Randle *et al.* 2009; Grossenbacher *et al.* 2015). In support of this argument, ecologists and evolutionary biologists have established that the distribution and range size of plant species are influenced strongly by climate (Parker 1963; Stephenson 1990; Park & Potter 2015), and large geographic range size is thought to be associated with wider niche breadths (Brown 1984; Slatyer *et al.* 2013). Species occurring at higher latitudes have also been hypothesized to have broader environmental tolerances due to larger seasonal fluctuations (Stevens, 1989; but see Šizling *et al.*, 2015). Additionally, selfing has been hypothesized to promote local adaptation and niche divergence by converting non-additive genetic variance to additive variance for tolerance to new habitats, thus facilitating expansion into novel climatic conditions (Lande 1977; Kirkpatrick 2000; Levin 2010).

An alternative interpretation is that the switch to self-fertilization is an evolutionary “dead-end” (Dobzhansky 1950; Stebbins 1957). Selfing is associated with increased homozygosity, reduced effective population sizes (Pollak 1987; Schoen & Brown 1991), increased accumulation of mutations (Heller & Smith 1978; Morran *et al.* 2009), and reduced genetic diversity (Jarne & Städler 1995; Hamrick & Godt 1996; Nybom 2004; Glemin *et al.* 2006). These effects are hypothesized to limit the ability of selfers to adapt to different environments and extend their ranges relative to their outcrossing relatives (Crawford & Whitney 2010). If selfing is an evolutionary dead-end, then selfers should not have greater niche breadths than their outcrossing sister taxa. Moreover, it is possible that geographic range and niche breadth are decoupled over short evolutionary times (Randle *et al.* 2009). Under this scenario, we would hypothesize that selfers may inhabit larger geographic ranges that exhibit relatively little climatic variation relative to their outcrossing sisters. Furthermore, we would also expect that the lack of genetic heterozygosity and adaptive potential limit the degree to which climatic niches of selfers species can diverge from each other, resulting in greater niche overlap between selfing sister species than between outcrossing sisters.

Here, we test this hypothesis by examining whether selfers with larger geographic ranges than their outcrossing relatives also have greater niche breadths, and how the relationship between range size and niche breadth changes over time. We also examine niche overlap between pairs of selfing sister taxa and between pairs of outcrossing sister taxa to determine whether niches are less likely to change after the shift to selfing. Niche overlap is negatively proportionate to the amount of niche change since divergence, and thus may reflect the potential of species to expand their niches (Broennimann *et al.* 2007; Turner *et al.* 2015). Finally, as the accumulation of deleterious mutations is thought to limit long-term adaptive potential and fitness in selfing lineages (Heller & Smith 1978), we explore whether rates of mutation accumulation differ significantly between selfers and outcrossers. We hypothesize that freedom from mate limitation initially allows selfers to expand their geographic ranges and niche breadth. However, as homozygosity and the accumulation of mildly deleterious mutations increase due to inbreeding, the adaptive potential of selfers should decrease more rapidly than that of outcrossers, resulting in more constrained niches.

## Materials and Methods

All analyses described below were done in R (R Core Team 2013); detailed information on the packages used are provided in Table S1.

### Dataset

To estimate and compare the niche breadth of selfing and outcrossing species, we used a previously published dataset that collated 54 studies describing plant species’ mating systems from 20 well supported, phylogenetically divergent clades representing 15 families (Grossenbacher *et al.* 2015). All taxa were included in previously published species-level phylogenies containing at least one predominantly selfing species and one predominantly outcrossing species. Species were classified as predominantly selfing when outcrossing rates were below 0.2 and predominantly outcrossing when above 0.8 (see Table S3 in Grossenbacher *et al.*, 2015). Species with outcrossing rates in between, or those exhibiting extensive among-population variation in outcrossing rates and traits associated with outcrossing were treated as having a variable mating system and not used in this study. We also used previously published time-calibrated phylogenies for all 20 clades based on internal transcribed spacer (*nrITS*) sequences (Grossenbacher *et al.* 2015, 2016). Sister species were identified in a subset of 9000 trees from the posterior distribution for each clade. The posterior probability of each sister pair, *i.e.*, the proportion of trees in which the two species were sister, was used as a measure of phylogenetic uncertainty. A total of 498 sister species pairs were identified, of which 194 differed in mating system.

### Estimating climatic niche breadth and overlap

We used curated geographic records (excluding those with coordinate accuracy > 100 km, coordinates failing to match the locality description, or those with taxonomic misidentifications), from the Global Biodiversity Information Facility (http://www.gbif.org) to infer the environmental conditions each species occupies (Grossenbacher *et al.* 2015, 2016). Among abiotic environmental variables, temperature (Parker 1963) and moisture (Stephenson 1990; Pigott & Pigott 1993) have been shown to strongly influence plant ranges (Holdridge 1947). We thus examined global data on 19 bioclimatic variables at 2.5 arc-minute resolution derived from monthly temperature and rainfall values (Nix 1986; Busby 1991; Hijmans *et al.* 2005). To maximize comparability with previous examinations of geographic range (Grossenbacher *et al.* 2015, 2016), we recorded the minimum, maximum, and standard deviations (SD) of each bioclimatic variable in species’ climatic ranges, which together define univariate climatic niche breadth (May & MacArthur 1972). We also examined each species’ niche breadth by calculating the average Euclidian distance between all points in each species’ range and the center of their distribution in in 19-dimensional climatic space (*i.e.*, multivariate standard deviation). As Euclidian distance measures can be sensitive to covariance among variables, we repeated this process on a subset of seven bioclimatic variables likely associated with the distribution of plant species, and whose pairwise Pearson’s correlation coefficient *r* < 0.75: isothermality (BIO2), minimum temperature of the coldest month (BIO6), mean temperature of the wettest quarter (BIO8), precipitation of wettest month (BIO13), precipitation seasonality (BIO15), precipitation of the warmest quarter (BIO18), and precipitation of the coldest quarter (BIO19). We also used PCA to summarize all the bioclimatic variables, and calculated niche breadth as the area of the convex hull surrounding each species’ points of occurrence in climate space defined by the first two principal axes. Species ranges were defined as the summed area of occupied grid cells across cell sizes of 0.05, 0.1, 0.5, or 1 decimal degree, corresponding roughly to 25, 100, 2500, and 10,000 km^2^ respectively (Grossenbacher *et al.* 2015, 2016). All subsequent analyses were done using niches derived from each of these cell sizes to assess whether the results were sensitive to the spatial grain of estimation.

To examine the degree of niche overlap between sister species, we used the approach developed by Broennimann *et al.* (2012), which has been shown to be robust to errors and biases associated with the estimation of niche overlap. This method compares the environmental conditions available for a species within a defined study extent with its observed occurrences and calculates the available environmental space defined by the first two principal axes. These extents again were delimited as occupied cells of 0.05, 0.1, 0.5, and 1 decimal degrees to account for different spatial grains. The same 19 bioclimatic variables used above were used for the multivariate PCA. Sampling bias was corrected by employing a Gaussian kernel density smoothing approach. The degree of niche overlap between each sister species pair was calculated using Schoener’s D (Schoener 1968) and modified Hellinger distance (I) (Warren *et al.* 2008), which vary from 0 (no niche overlap) to 1 (identical niche).

### Statistical analyses

Linear mixed effects models were used to test differences in *ln*-transformed niche breadth between outcrossing and selfing sister species. Mating system was treated as a fixed effect. Genus (or section, in the case of *Oenothera*) and sister-pair identity entered the model as random factors, and we estimated slopes for each genus. The posterior probability of each sister species pair was included as a weighting factor in our models to account for phylogenetic uncertainty. We included the interaction of divergence time (*ln*-transformed) with mating system as a fixed effect in the model to test whether the effect of divergence time on niche breadth co-varied with mating system. This test also included a random effect for genus-specific mating system. As niche breadth may be affected by ploidy and lifespan (Morishima *et al.* 1984; Thompson *et al.* 2014), we also ran these analyses including only sister pairs that did not differ in by these potentially correlated traits.

Niche overlap values range from 0 to 1. These bounds, and the right-skewed distributions of these measures, violate the assumptions of standard linear models (Ramalho *et al.* 2011). Hence, we used fractional logit regression models (Papke & Wooldridge 1996) to test whether the mating systems of sister-species pairs (s-s, s-o, o-o) influenced niche overlap, with sister-pair mating system as a categorical predictor and niche overlap (Schoener’s D or Hellinger’s I) as the response variable. The posterior probability of each sister-species pair again was included as a weighting factor to account for phylogenetic uncertainty, and models were fit using maximum likelihood. We similarly used fractional logit models to test whether time since divergence (*ln*-transformed) influenced niche overlap. All analyses were replicated across the four spatial scales defined above to examine whether our results were robust to the spatial scale at which species niche was estimated. Finally, to address the possibility that certain clades may heavily influence the overall results, we ran these analyses dropping each individual genus alternatively. Only those cases for which dropping a genus (or section in *Oenothera*) affected the statistical significance of the analyses are reported.

Last, we used linear mixed models to test whether the rate at which lineages accumulate mutations was influenced by mating system, and how this relationship changes over time. The interaction of divergence time (*ln*-transformed) with mating system entered the model as a fixed effect, and we estimated a genus-specific random slope for divergence time. We restricted these analyses to the 42 sister pairs with annual life histories and identical ploidy levels because both of these factors can influence mutation fixation rates. Annuals have higher rates of molecular evolution than perennials, and unlike perennials display rates similar to other closely-related annuals (Andreasen & Baldwin 2001). Mutation accumulation rates were calculated by dividing the number of nucleotide substitutions since divergence by divergence time for each selfer-outcrosser sister-species pair across all phylogenies (Smith & Donoghue 2008). In all cases, we report pseudo-R-squared values as measures of the variation explained by fixed and random effects.

## Results

Mating system did not significantly influence niche breadth measured as multivariate climate space (Fig. 1A, Table 1) or based on individual bioclimatic variables (p > 0.05 all cases; Table S2). The relationship between mating system and niche breadth was inconsistent, in contrast to that between mating system and geographic range size. Comparisons of niche breadth for the uncorrelated subset of bioclimatic variables yielded similar results, as did comparisons based on the first two principal axes of climate space occupied by species (PCA; p > 0.05; Fig. S1, Table S3). These results were consistent across all spatial grains and when excluding sister pairs that differed in ploidy and life history (Table S4, S5). However, the niche breadth of selfing species tended to decrease more over evolutionary time than the niche breadth of their outcrossing sisters (divergence time × mating system: p < 0.05; Fig. 2A, Table S6).

**Figure 1.**
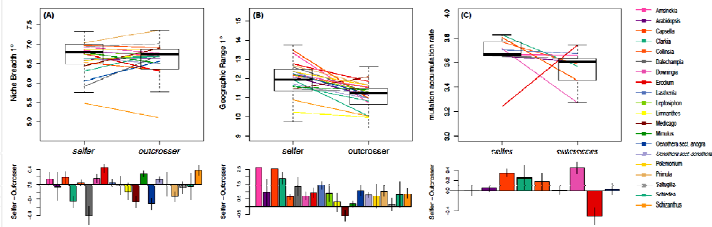
Comparisons of niche breadth, geographic range, and mutation accumulation rates among selfers and outcrossers. (A) Top panel: Box plots of predicted (fitted) niche breadth of selfing and outcrossing sister species assessed in 19 dimensional climate space and at 1° resolution. Colored line segments indicate predicted slopes for each of 20 clades and the vertical axis is natural logarithmic scale. Bottom bar charts: average sister species log difference in niche breadth for each of 20 clades, with vertical lines representing standard errors. (B) Top panel: Box plots of predicted range size of selfing and outcrossing sister species assessed at 1° resolution. Bottom bar charts: average sister-species log difference in range size, with vertical lines representing standard errors. (C) Top panel: Box plots of predicted rate of mutation accumulation of annual selfing and outcrossing sister species. Bottom bar charts: average sister-species log difference in rate, with vertical lines representing standard errors. Clade averages are only used for illustration purposes, and statistical analyses were performed with individual species-pair estimates.

**Figure 2.**
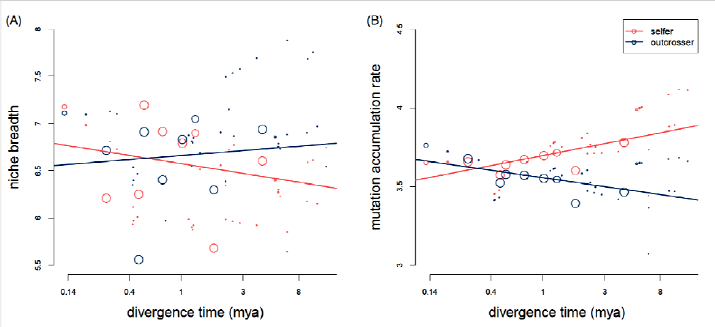
Niche breadth (A) and mutation accumulation rate (B) as a function of divergence time for selfing and outcrossing sister species at 1° resolution. The size of the circles represent the posterior probability of which the focal species pair were each other’s sister taxon. The line segments represent the linear regression results for selfers (pink) and outcrossers (blue). Niche breadth is log-transformed and divergence time is on a back-transformed natural logarithmic scale.

**Table 1.** Results of five separate linear mixed models analyzing the effect of mating system on species’ niche breadth in 19 dimensional climate space estimated at four different spatial resolutions. The categorical coefficient estimates are log-odds ratios and represent departures from the “outcrosser” mating category. Marg. R^2^ represents the proportion of variance in explained by mating system and Cond. R^2^ values are the variance explained by the entire model.

| Resolution | Estimate | Std. Error | df | t | p-value | Marg. $R^2$ | Cond. $R^2$ |
| --- | --- | --- | --- | --- | --- | --- | --- |
| *Response: ln-transformed niche breadth* |  |  |  |  |  |  |  |
| 0.05° | -0.11 | 0.24 | 18.28 | -0.49 | 0.63 | 0.00 | 0.97 |
| 0.1° | -0.17 | 0.19 | 16.28 | -0.88 | 0.39 | 0.01 | 0.95 |
| 0.5° | -0.06 | 0.14 | 16.63 | -0.41 | 0.69 | 0.00 | 0.95 |
| 1° | -0.05 | 0.10 | 14.50 | -0.53 | 0.60 | 0.00 | 0.95 |

To compare niche expansion ability between selfers and outcrossers, we assessed the proportion of shared niche space (overlap) among sister taxa of different mating system combinations: selfing-selfing (s-s), selfing-outcrossing (s-o), and outcrossing-outcrossing (o-o). Patterns of niche overlap were significantly influenced by mating system. In particular, selfing sister pairs (s-s) had significantly greater degrees of niche overlap (e.g., Schoener’s D = 0.23) than sister pairs with at least one outcrossing species (e.g., s-o: D=0.15; o-o: D=0.17) regardless of the metric examined (p < 0.05; Fig. 3; Table S7). These patterns were robust to the removal of all genera except *Medicago* (p > 0.05; Table S8). Although the distribution of divergence times differed among the three mating system pairs (Grossenbacher *et al.* 2016), niche overlap was not influenced by divergence time (Table S9).

**Figure 3.**
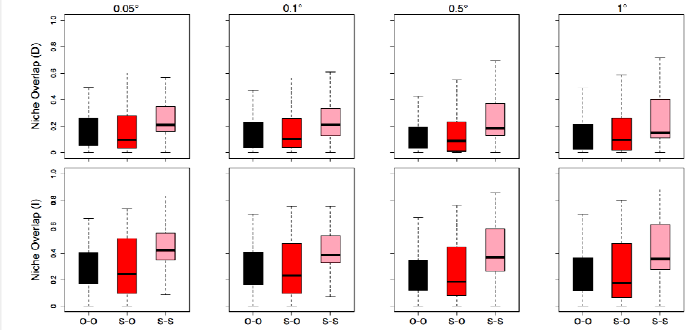
Boxplots of two metrics of sister pair niche overlap by mating system category at four spatial resolutions; outcrosser–outcrosser (s-s, dark grey), selfer–outcrosser (o-o, red), selfer–selfer (s-o, pink).

Finally, to investigate whether genetic degradation might limit selfers’ niche expansion and differentiation over time, we examined mutation accumulation rates across sister taxa with different mating systems. Wide variation was observed in the mutation accumulation rates among the taxa we examined. Selfers had higher mutation accumulation rates than their outcrossing sisters in general (Fig. 1C). Together with time since divergence, the shift to selfing explained a significant proportion of this variation (R^2^ > 0.27; Table S10). The rate of mutation accumulation was predicted to increase through time for selfers while concomitantly decreasing for their outcrossing sisters (Fig. 2B).

## Discussion

Even though selfers occupy larger geographic ranges and higher latitudes than their outcrossing sisters (Grossenbacher *et al.* 2015), we find that this does not necessarily translate into greater climatic niche breadth. Our results indicate that niche breadth and range size are decoupled, potentially leading to species with large geographic ranges but narrow climatic niches. For three reasons discussed below, we argue that the large range sizes currently observed for selfing plant species are an evolutionarily transient phenomenon.

### Mating system does not consistently predict niche breadth

Our niche breadth analyses do not support a tight association between range size and niche breadth. Neither do our results support the hypothesis that selfers should have wider niche breadth than
their outcrossing sisters due to their higher latitude ranges. This suggests that although the reproductive assurance offered by self-fertilization may allow selfers to expand their geographic range (Grossenbacher *et al.* 2015), this expansion does not always occur into novel climates. Because the vagaries of geography can result in certain climates occurring more frequently than others, species adapted to common climatic conditions may exhibit larger geographic distributions than their climatic niches would suggest (Burgman 1989; Hanski *et al.* 1993; Gaston & Spicer 2001; Thompson & Ceriani 2003). For instance, the outcrossing *Medicago edgeworthii* Sirj. grows in a wider range of climates than its close selfing relative *M.radiata* L. despite having a much more restricted geographic range centered at a lower latitude (Fig. 4). Furthermore, given the penchant of selfers to exist in comparative low abundance at small fragmented habitats, their geographic ranges; and the climatic space they occupy; may have been overestimated relative to their outcrossing relatives.

**Figure 4.**
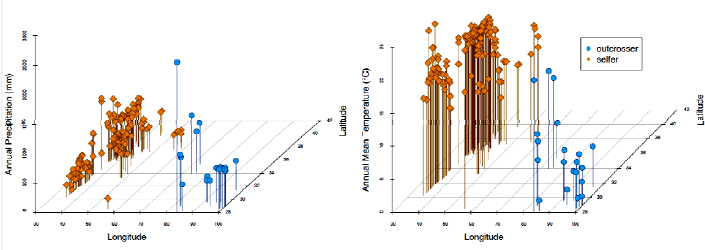
Geographic range and climatic range inhabited by *Medicago edgeworthii* (outcrosser) and *Medicago radiata* (selfer). Points represent the geographic locations of each species.

### Niche divergence is negatively impacted by selfing

It has been suggested that selfing species have a reduced capacity to adapt to different environments (Crow 1992; Morran *et al.* 2009). This lack of adaptability was supported by our finding that the degree of niche overlap was higher among selfing sister-species pairs than outcrossing pairs. This result suggests that species’ niches are slow to diverge following the transition to selfing. As there are no known instances of outcrossing revolving from selfing lineages, we can reasonably assume that selfing sister species diverged post-transition (i.e., their most recent common ancestor is likely a selfer). Assuming a degree of spatial autocorrelation among environmental conditions, this result seemingly supports the long-held theory that autonomous self-fertilization facilitates range overlap of closely-related species (Antonovics & Bradshaw 1970). Numerous mechanisms can promote the coexistence of selfers and their close relatives, including minor range shifts following peri- or parapatric speciation that promote early secondary range overlap, reproductive isolation from ancestral or sister species, and reduced competition for pollinators (Grossenbacher *et al.* 2016). However, the higher level of niche overlap that we observed for selfing sister-species pairs did not result from geographic proximity, as there was no association between mating system and co-occurrence among sister species (Grossenbacher *et al.* 2016). This suggests that once species have transitioned to selfing, they may be unable to establish in new climates as readily as their outcrossing relatives, and thus expand their geographic range by colonizing familiar environments.

Finally, although the effect of selfing pairs was no longer significant when the genus *Medicago* was removed from our analysis (Table S8; see also ref. 25), this result is likely a consequence of a substantial reduction in sample size. *Medicago* includes 39 selfing sister pairs: over 43% of the weighted sample of selfing sister pairs and more than any other genus included in our study. Nonetheless, these results suggest that climatic niche divergence is not facilitated by selfing. Indeed, in Gallagher *et al.*’s (2010) examination of niche shifts in 26 plant species introduced to Australia, the six species that do not exhibit evidence of niche shifts are all primarily self-pollinating. Furthermore, a number of recent studies have illustrated the greater potential for niche expansion by outcrossing species (Broennimann *et al.* 2007; Petitpierre *et al.* 2012; Gallien *et al.* 2016).

### Selfing leads to decreased niche breadth over time

Among the species we examined, selfers did not uniformly have smaller niches than their outcrossing sisters, but the climatic ranges of the former were predicted to decrease significantly more rapidly over time. Thus, the climatic niches of selfing species eventually will become narrower than the niches of related outcrossing species, irrespective of their initial niche breadth. It is possible that the decrease of a selfer’s niche over time can be attributed to genetic impoverishment caused by inbreeding. The reduction in effective population size that accompanies selfing limits both positive and purifying selection, increases the fixation of deleterious mutations, and impairs the ability of a species to adapt to novel conditions over time (Bachtrog & Charlesworth 2002; Wright & Andolfatto 2008).

Along these lines, significantly higher amounts of nucleotide substitutions were fixed in selfing species following divergence from their outcrossing sisters. As molecular evolutionary rates are similar among closely-related annual plant species (Andreasen & Baldwin 2001), the differences in branch lengths (*i.e.*, substitutions) we observed between sefling and outcrossing sisters reflect the faster rates at which mutations were **fixed** in selfing lineages. Although a non-coding region was used to build the phylogenetic trees in our dataset, our results demonstrate that selfers accumulate non-lethal mutations more rapidly. This reflects the lack of heterozygosity present in selfer genomes, and small effective population sizes As with selectively neutral mutations, mildly deleterious, non-lethal mutations can also escape purifying selection, and thus are likely to more rapidly accumulate in selfing lineages (Slotte *et al.* 2013). The comparatively small sizes of selfing species’ genomes increases the chance of deleterious mutations being fixed via linked (background) selection as well (Hudson & Kaplan 1995; Albach & Greilhuber 2004; Charlesworth 2012). Our results further suggest that mutation accumulation rates of selfers will only increase through time. As each generation of selfing reduces heterozygosity by 50%, the genomes of primarily selfing species will rapidly reach near clonal status. Similarly, selfing is expected to reduce effective population size by a factor of two, or more due to a high probability of experiencing bottlenecks through founding effects and a more pronounced effects of linked selection (Jarne 1995; Charlesworth & Wright 2001; Hartfield 2016). In this case, non-lethal deleterious mutations will become fixed almost as soon as they arise (Glémin & Ronfort 2013). Thus, lineages of selfing species may persist for less time than outcrossing lineages, and extant selfing lineages have accordingly been found to be evolutionarily recent (Foxe *et al.* 2009; Escobar *et al.* 2010; Ness *et al.* 2010; Busch *et al.* 2011; Pettengill & Moeller 2012).

Additional factors may influence the rate at which mutations are fixed or niche breadth changes, but we minimized the potential effects of unaccounted variables by comparing closely-related sister species. Although our analyses are limited in scope and do not enable us to make a direct or causal connection between the predicted decrease in niche breadth of selfing species and long-term costs of reduced genetic diversity, previous studies have shown them to be linked (Noy *et al.* 1987; Morran *et al.* 2009), and selfing lineages have been shown to experience considerable accumulation of deleterious mutations over relatively short timescales (Hu *et al.* 2011; Slotte *et al.* 2013).

### Reconciling geographic range and climatic niche breadth

In general, species with higher levels of genetic diversity should maintain populations across greater environmental heterogeneity, thus facilitating larger geographic ranges than species with narrow ecological niches (Brown 1984). However, it is possible that the realized niche of selfing species is closer to their fundamental niche than it is for their outcrossing sisters. The fundamental niche refers to the environmental requirements of a species to maintain a population indefinitely, independent of species interactions and immigration (Hutchinson 1957; Holt 2009). The realized niche is the proportion of a species’ fundamental niche that remains after accounting for interspecific interactions (e.g., competition), dispersal limitation, and the lack of suitable contemporary environments (Colwell & Rangel 2009). The release from mate limitation allows selfers to colonize small or fragmented habitats, enabling them to explore a larger extent of their fundamental niche (i.e., increase of realized niche). The initial purging of recessive deleterious alleles could also contribute to the expansion of selfers’ realized niche (Peterson & Kay 2015). Such a scenario could translate to a transiently larger species range for selfers, but the limitations of an increasingly homogeneous genome should become apparent over time. Mutations accumulate as heterozygosity decreases, the fundamental niche contracts, and eventually the geographic range shrinks (Fig. 5).

**Figure 5.**
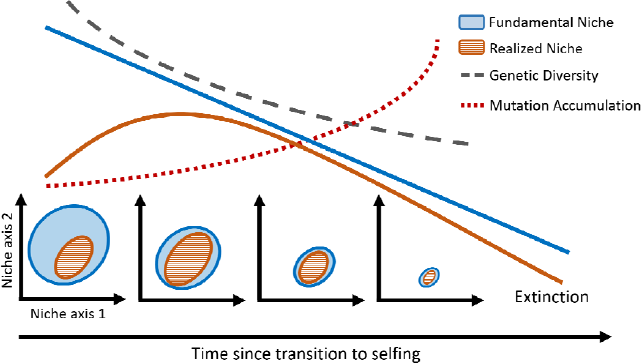
Expected changes in genetic diversity, mutation accumulation, fundamental niche, and realized niche following transition to selfing.

Like previous studies (Grossenbacher *et al.* 2015, 2016), our analyses are correlative and we cannot demonstrate a causal relationship between niche breadth and mating system. We restricted our analysis to definitively selfing or outcrossing species, but multiple mechanisms affect intermediate levels of self-fertilization, even within populations (Goodwillie *et al.* 2005). Despite the inclusion of the largely tropical genera *Dalechampia* and *Schiedea*, most of the clades we analyzed have primarily temperate distributions, and different patterns might have been observed if more tropical taxa had been included. As more data become available, we can examine further the relationships between the degree of selfing, range size, and niche breadth that account for additional factors, including biotic interactions.

Nonetheless, our observations that, relative to outcrossing sister taxa, the climatic niches of selfing species are slower to differentiate (Fig. 3) and stand to become narrower over time with genetic degradation (Fig 2) are consistent with the well-established idea that selfing is an evolutionary dead-end (Dobzhansky 1950; Stebbins 1957). Despite high transition rates to selfing, < 15% of extant seed plants are predominantly selfing (Goodwillie *et al.* 2005; Igic & Kohn 2006) and transitions from selfing to outcrossing occur rarely, if ever (Igic & Busch 2013). Furthermore, selfing lineages have been shown to be younger than outcrossing ones, implying that they are more short-lived (Holsinger 2000). Genetic impoverishment and accumulation of mildly deleterious alleles may not manifest as short-term losses of fitness or geographic range in all selfing species, but it likely will affect their potential for evolutionary adaptation (Honnay & Jacquemyn 2007) and eventually outweigh any (initial) advantages of self-fertilization (Fig. 5). Indeed, simulations have demonstrated that the greater genetic load in self-compatible lineages results in overall increases in time to adaptation and extinction risks regardless of self-fertilization rates (Peterson & Kay 2015). Given rapid rates of recent climatic change, this may have severe consequences in the near future, especially since many plant species lack sufficient ability to track the shifting climate northward or upward (Honnay *et al.* 2002). The larger geographic range and comparable niche breadth of many selfers most likely is a temporary phenomenon caused by an expanded realized niche, and may be a snapshot of the early stages of a temporal spiral towards extinction.

## Acknowledgements

The authors thank D. Grossenbacher, R. D. Briscoe Runquist, E. E. Goldberg, and Y. Brandvain for making their data and analyses available, S. Worthington for statistical advice, and D. S. Barrington, C. G. Willis, O. Razafindratsima, and T. J. Davies for their insightful comments on the project and manuscript. We are also grateful for the invaluable feedback from the editor and anonymous reviewers. This work was made possible by the Harvard University Herbaria. AME’s participation in this project was supported by the Harvard Forest.

## Data Accessibility

Phylogenetic trees, occurrence data, and mating 526 system data are available at: Dryad doi:10.5061/dryad.hv117

## Author Contributions

D.S.P conceived the study idea and performed all analyses; A.M.E. advised statistical analyses; D.S.P., A.M.E., and C.C.D. interpreted the results; and D.S.P and C.C.D. wrote the manuscript. All authors contributed significantly to revisions.

## Supporting information

Figure S1: Comparisons of niche breadth and geographic range among selfers and outcrossers.

Table S1: R Packages used in analyses.

Table S2: Results of linear mixed models analyzing the effect of mating system on species’ range (max – min) and standard deviation (SD) across 19 bioclimatic variables estimated at four different spatial resolutions.

Table S3: Results of five separate linear mixed models analyzing the effect of mating system on species’ niche breadth in 7 dimensional climate space estimated at four different spatial resolutions.

Table S4: Results of ten separate linear mixed models analyzing the effect of mating system on species’ niche breadth in 19 dimensional climate space estimated at four different spatial resolutions.

Table S5: Results of linear mixed models analyzing the effect of mating system on species’ range (max – min) and standard deviation (SD) across 19 bioclimatic variables estimated at four different spatial resolutions.

Table S6: Results of four separate linear mixed models examining the effect of mating system, divergence ce time, and their interaction on species’ niche breadth assessed across four spatial resolutions.

Table S7: Results of fractional logit regression models assessing the effect of sister pair mating system on niche overlap at four spatial scales.

Table S8: Results of fractional logit regression models assessing the effect of sister pair mating system on niche overlap at four spatial scales, sequentially removing each genus.

Table S9: Results of fractional logit regression models assessing the effect of sister pair divergence time on niche overlap four spatial scales. The coefficient estimates are log-odds ratios.

Table S10: Results of a linear mixed model examining the effect of mating system, divergence time, and their interaction on species’ rates of molecular evolution.

